# Deep learning based reconstruction of embryonic cell-division cycle from label-free microscopy time-series of evolutionarily diverse nematodes

**DOI:** 10.1101/2024.05.09.593369

**Authors:** Dhruv Khatri, Chaitanya A. Athale

## Abstract

Microscopy of cellular dynamics during embryogenesis of non-model organisms can be technically challenging due to limitations of molecular labelling methods. Label-free differential interference contrast (DIC) microscopy of the first embryonic cell division of nematodes related to *Caenorhabditis elegans* has been successfully employed to examine the constraints and divergence of intra-cellular mechanisms during this asymmetric cell division. However, identifying stages of the cell division cycle were performed interactively, pointing to a need to automate of cell stage identification from DIC microscopy. To this end, we have trained deep convolutional neural networks (CNNs), both pre-existing such as ResNet, VGGNet and EfficientNet, and a customized shallow network, EvoCellNet, to automatically classify first-embryonic division into the stages: (i) pro-nuclear migration and (ii) centration and rotation, (iii) spindle elongation and (iv) cytokinesis, with all networks performing with 91% or greater accuracy. The activations of the networks superimposed on the images result in segmentation-free detection of intracellular features such as pro-nuclei, spindle and spindle-poles in case of the shallow EvoCellNet, while ResNet, VGGNet and and EfficientNet detect large-scale, features that are less biologically meaningful. The UMAP space representation combined with support vector machines (SVM) allows for stage boundary identification and recovers a cyclical map connecting the states (i) to (iv) of the division. This approach could be used to automate quantification of cell division stages and sub-cellular dynamics without explicit labelling in label-free microscopy.

**Summary:** We have trained multiple convolutional neural networks (CNNs) to classify the stages of cell division from the first embryonic division of diverse nematodes, evolutionarily related to *Caenorhabditis elegans*. We find two classifiers, VggNet and a customized EvoCellNet, can detect intracellular features and a UMAP representation can reconstruct the cyclical progression of first embryonic division from related species.

## Introduction

The nematode *Caenorhabditis elegans* has formed a model for developmental biology, genetics and cell biology due to its short generation time, transparency of the early stages and ease of genetic manipulation [1]. In particular, this organism is well-suited for microscopy of the early stages of development due to its transparent nature and presence of intracellular structures that provide image contrast, even in the absence of molecular labels [2, 3]. At a sub-cellular scale this has been exploited to examine intracellular dynamics. First embryonic division serves as a model for both early embryonic division as well as the basic tenets of asymmetric division. In *C. elegans* it begins with the entry of sperm into the oocyte [4], that establishes anterior-posterior (AP) polarity and triggers the completion of oocyte meiosis [5]. The subsequent step involves the convergence of the two pronuclei towards each other, followed by their fusion to form a centrosome-pronucleus complex [4]. This complex moves to the center of the embryo and aligns itself along the AP axis, leading to the formation of a mitotic spindle that separates the chromosomes into two opposite poles. The progression to anaphase is associated in *C. elegans* with a characteristic oscillatory movement that has been identified as resulting from asymmetric pulling forces acting on astral microtubules (MTs) exerted by dynein motors anchored at the cell cortex in anterior and posterior poles, with force asymmetry and MT dynamics driving oscillations [6, 7, 8]. Eventually a cytokinetic furrow is formed, which divides the cytoplasm, giving rise to two asymmetric daughter cells. The cytoplasm fluid mechanical properties have been characterized to result in an effective viscosity ranging between 0.6 to 1 Pa-s based on either microrheology of tracking passive yolk granule movement [6, 9, 10], single particle tracking of injected fluorescent sub-micron particles [11] and spindle displacement and recoil using magnetic tweezers [12]. AAcross-speciesross-species, some closer and some evolutionarily more distant to *C. elegans* have demonstrated spindle movements in Anaphase vary with very little correlation to phylogenetic distance [13]. In contrast the mechanics of the cortical pulling force (F) acting on the spindle, effective cytoplasmic viscosity (*η_eff_*) and embryo size (*L*) when combined as *ω* = *F/*(*L· η_eff_*) is predictive of spindle oscillation onset [10]. Comparative studies on spindle dynamics have revealed universal scaling relationships between embryo size and spindle dynamics in *C. elegans* strains alone [14]. However, to look at the scaling of physical properties and cell division dynamics in more diverse species, we would first need a way to classify the stages of cell division during first embryonic division without the use of molecular markers. This is particularly relevant since outside of *C. elegans* the genetic modifications have proved technically challenging.

The analysis of DIC microscopy for sub-cellular morphology in nematode embryos is feasible due to the transparent nature of the cells, with pronuclei, centrosome-pronuclear complex, and mitotic spindle visible to the eye forming a region of uniform intensity, surrounded by the granular cytoplasm. This has been exploited to examine spindle dynamics using a semi-automated active contours approach to automate the measurement of spindle dynamics [15]. Additionally the distinction between organelles and the cytoplasm is used to identify mutants in functional genomic screens [16, 17, 18, 19]. Additionally, automated analysis has also been developed to study embryo shape [20], pronuclear migration during fertilization [21] and cytoplasmic yolk granule mobility [9]. However, these tools require a cell stage identification that is either done at acquisition or selection of those frame. A general tool to combine intracellular dynamics with an input of the whole cell cycle image time-series could be used for larger screening projects in future. Despite the well-defined stages of the cell division process in *C. elegans* and other nematode species, accurately measuring the duration of each phase is a time-consuming and laborious task. Manual annotation of time-lapse microscopy images requires expertise and can be prone to human error for large datasets. Therefore, there is a need for automated tools to accurately and efficiently annotate cell division time scales across diverse nematode embryos. This will not only allow for rapid quantification cell stage differences but also to related sub-cellular processes which regulate cell stage progression.

Recently, convolutional neural networks (CNNs) have been applied in different areas of biology such as cancer classification [22], cell cycle analysis [23, 24] and cell phenotyping [25, 26]. In particular, *DeepCycle* demonstrates the ability of CNNs to quantify cell stage progression in fluorescent cells. However, deep learning (DL) tools cannot be directly applied to novel datasets without considerable re-training and optimization. Thus a tool to examine early embryonic cell cycle could be of some use to the field.

In this study, we have developed a convolutional neural network (CNN) to accurately classify morphologically distinct stages of the first embryonic division across various *Caenorhabditis* species. The average F1-score for classification of all species is 95%. The CNN features, visualized using Gradient-weighted Class Activation Mapping, Grad-CAM [27] correspond to stage-dependent structures that were identified across the *Caenorhabditis* genera. The method can be applied to label-free DIC microscopy data with no tuning parameters. To our knowledge, this is the first report on the application of CNN in classifying cell stages in label-free DIC microscopy data of diverse *Caenorhabditis* embryos.

## Results and Discussion

### Training CNNs to detect stages of first embryonic cell divisions of nematodes in DIC microscopy

Nematode embryos especially those of *Caenorhabditis elegans* are well suited for differential interference contrast (DIC) microscopy due to the transparent nature of the oocytes and the presence of a high density of birefringent yolk granules that provide contrast to spindle dynamics. The first embryonic division of 21 nematode species, evolutionarily related to *C. elegans*, had been recorded previously undergoing first-embryonic division [13]. This dataset was used develop a deep learning (DL) pipeline to automate the identification of the stages of fertilization and cell division. The time-series begin with the stage when male and female pro-nuclei are visible and end after completion of the first cell division as identified by the ingression of the cytokinetic furrow. Therefore, an annotation scheme consisting of four stages based on morphologically identifiable stages was created: (i) pro-nuclear migration (PM), (ii) nuclear centration and rotation (CR), (iii) spindle displacement (SD) and (iv) cytokinesis (Cy) (Figure 1A). Each time series was manually classified in a frame-by-frame manner into one of these classes, as seen in a representative montage of embryos of 5 related species namely *C. remanei*, *C. elegans*, *C. briggsae*, *C. angaria* and *C. portoensis* (Figure 1B). Qualitatively the points of transition appear to have similar morphological features, and we aimed to train the network to identify these.

**Figure 1:**
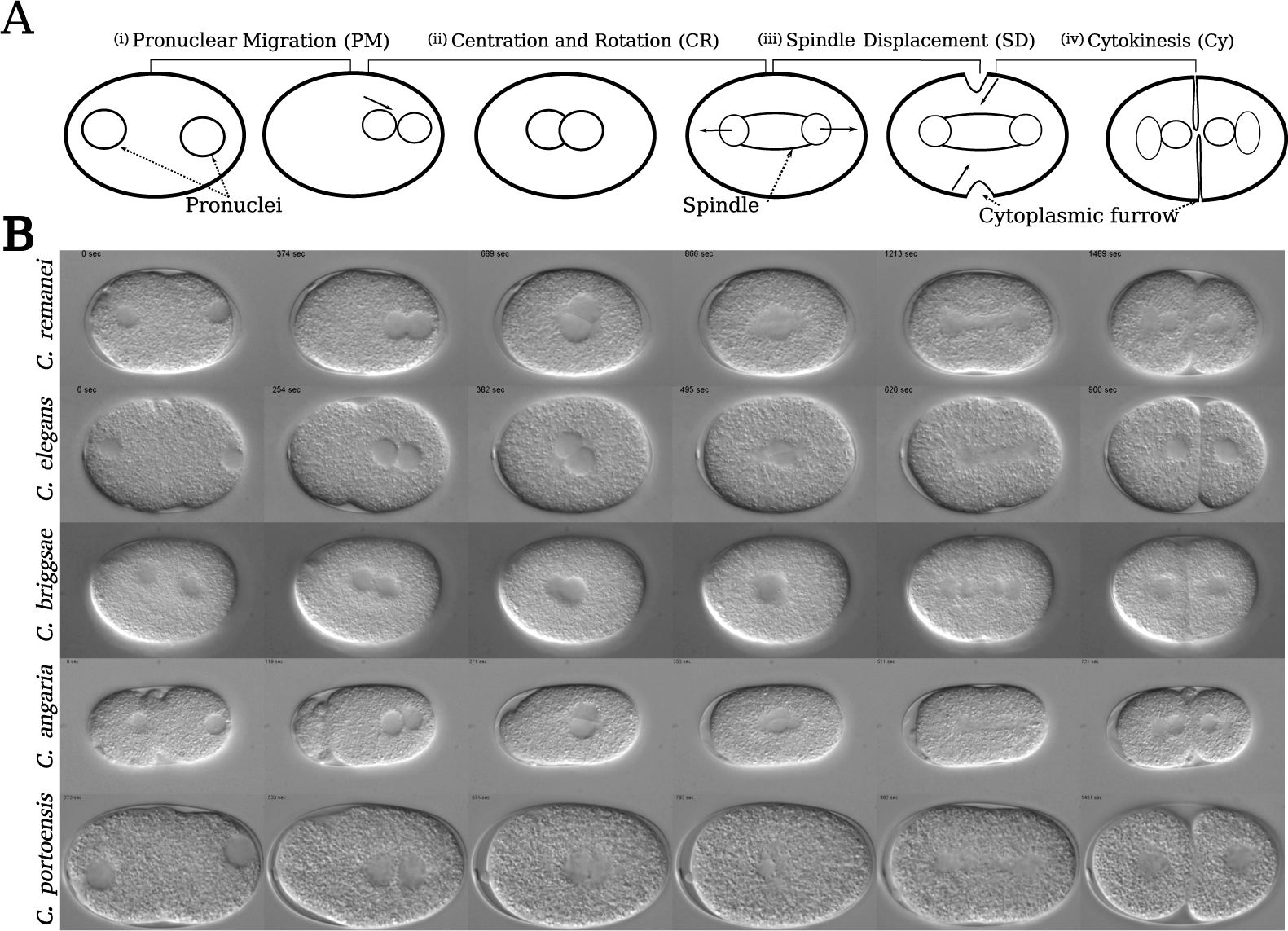
Stereotypical patterns during the first embryonic division across diverse nematode species. (A) A schematic of the first mitotic division in a typical nematode embryo is shown. We classify a given time-series of first embryonic division into four stages of (i) pronuclear migration (PM), (ii) centration and rotation (CR), (iii) spindle displacement (SD) and (iv) cytokinesis (Cy). Annotation of these stages is done by identifying sub-cellular events of pronuclear migration, spindle elongation and furrow indentation (indicated by bold arrows). (B) A montage of five *Caenorhabditis sp.* undergoing first embryonic division is shown. Species names is written at the start of every row. All DIC images taken from a published dataset [13]. The montage visualizes stereotypical morphology observed in a given stage across different species. These cytoplasmic features are formed by the two pronucleus and the mitotic spindle, which are visible as uniform intensity regions surrounded by a granular cytoplasm and a cell membrane.

We trained four CNN models to classify individual frames from time-series using Python code based on the PyTorch library (https://pytorch.org) ver. 2.1.0+cu118 [28]. Three of the networks used are considered state of the art for pattern recognition in images: deep residual network or ResNet [29], visual geometry group network or VggNet [30], and EfficientNet [31]. In addition, a shallow CNN that we refer to as EvoCellNet was also developed, in order to compare the effect of depth of the network and generality of previously developed architectures in detection. Based on the reported trade-off between complexity, accuracy and training time, the shallow network expected to take the least time to train. EvoCellNet is also the simplest of those tested, in terms of number of parameters, convolutional and linear layers (Table 1). Input data was augmented to reduce training bias with oversampling by affine transformations such as rotation, flipping in x- and y-axes and shearing which serves to make training independent of simple transformations and imaging conditions (Figure 2A). The training data consisted of 169 time-series, with the number of frames in each stage for the original training dataset different, with PM = 43,821, CR = 97,715, SD = 53,197 and CY = 63,336 due to the duration of the stages of first embryonic division (Figure 2B). For one epoch the number of images seen by the network is equalized by copying the input images, through image augmentation. Such oversampling of the classes was achieved by image augmentation. Networks were training was achieved by employing a stratified K-fold cross-validation split strategy where *k*=4, based on species and cell stage identity (Figure 2C). All models were trained using the same hyperparameters for comparison, and model weights were saved based on the minimum validation loss criterion.

**Figure 2:**
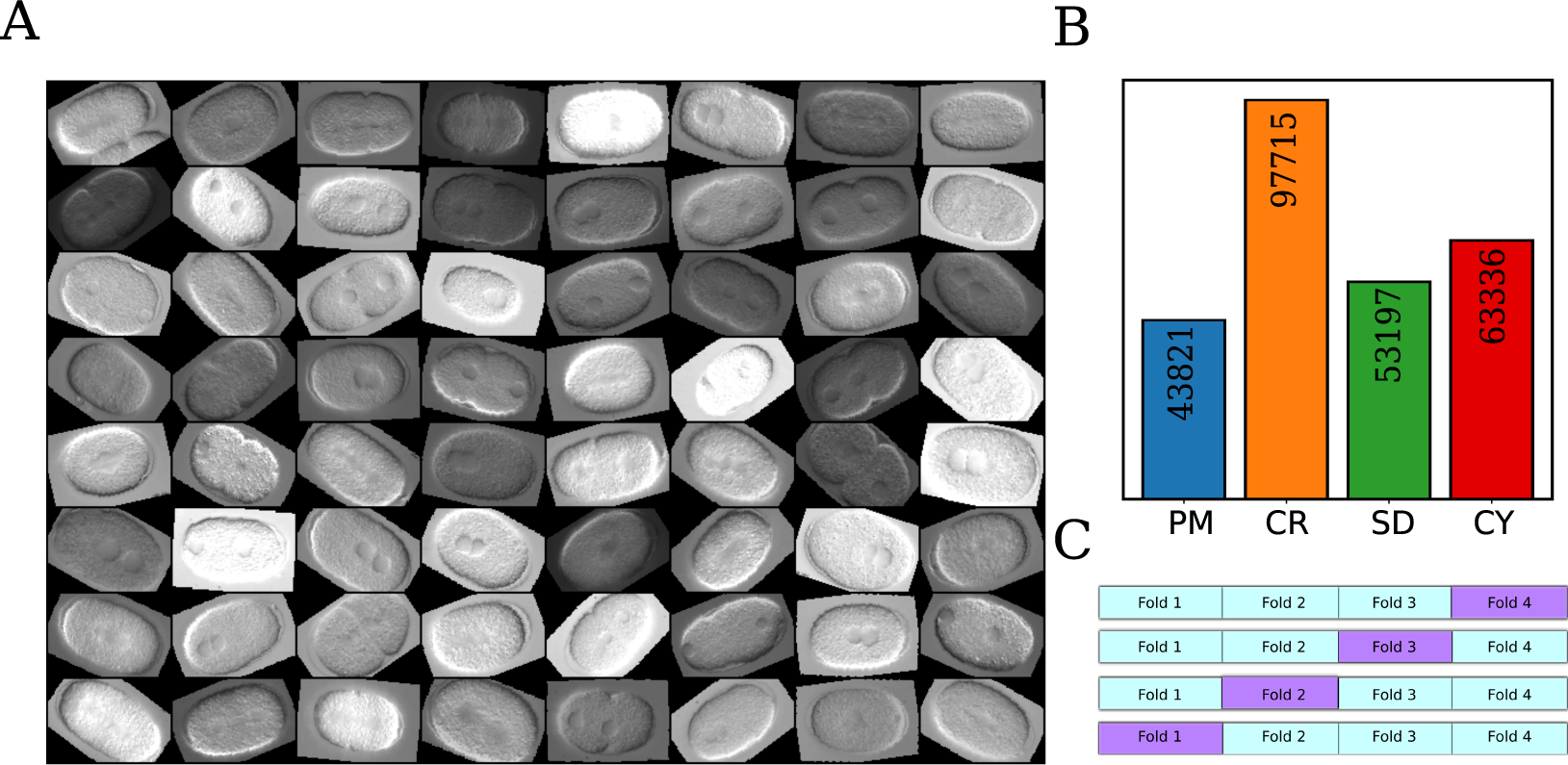
Training dataset with image augmentation. **(A)** The image montage consists of a batch of images (64) taken from the training dataset. Different augmentations such as image rotation, flipping (vertical and horizontal) and affine transformations have been applied randomly. **(B)** The bar chart shows the total number of frames obtained after annotation for each class. Color indicate annotated stages; PM (blue), CR (orange), SD (green) and CY (red). **(C)** The schematic represents k-fold cross validation scheme using which the training dataset was divided into four equal proportions maintaining equal proportion (25%) of each class.

**Table 1:** Configurational complexity of CNN models. The complexity of the previously de- scribed CNNs are compared in terms of number of parameters and layers both convolutional and linear.

| Network | Parameters ($\times 10^6$) | Convolutional layers | Linear layers |
| --- | --- | --- | --- |
| EvoCellNet | 0.7 | 7 | 0 |
| EfficientNet | 11 | 237 | 1 |
| ResNet | 11 | 19 | 1 |
| VggNet | 134 | 13 | 3 |

### Accuracy and loss of EvoCellNet in training is comparable to more complex networks

Our custom build CNN EvoCellNet has 6 convolutional layers followed by a one-dimensional convolutional layer with four output filters, with a 3x3 kernel size except for the first layer with a 5x5 kernel size (Figure 3A). The network architecture was optimized by trial-and-error. The input to the first layer is a 2D single channel image, an array of 256 x 256 pixels (height x width). The network uses skip connections, layer concatenation and max pooling to obtain the same size of all feature maps. Random removal of nodes during training with a probability *p* = 0.3, i.e. probabilistic dropout, was added after each layer except first layer, with the intention of generalization and avoiding over-fitting of the model to the data. The output from the last layer is passed through global average pooling layer producing a (N x C x 1 x 1) array, where N is batch size and C is the number of classes, and in total has 787,636 parameters. The input to the layers is normalized using batch normalization and the output from each layer is passed through a rectified linear units (ReLU) activation [32]. The training accuracy of all four networks resulting from average four-fold cross-validation, reaches the maximum in *∼*5 epochs remaining unchanged in 20 epochs in terms of both accuracy (Figure 3B) and loss (Figure 3C). There is little improvement if training is extended up to 50 epochs (Figure 3B,C *(inset)*). EvoCellNet compared to the deeper, more complex network architectures, shows comparable scores in terms of accuracy and loss during training (Figure S1A), but in validation has marginally lower scores (Figure S1B). The individual curves from each fold cross-validation show small differences in smoothness. The final convergence values of accuracy on the validation dataset are however similar, with EvoCellNet achieving an accuracy of 95%, compared to 97% for VggNet, 96% for EfficientNet, and 97% for ResNet. A comparison of precision, recall and F1 scores show all four networks perform very similarly to each other (Figure 4A, Table 2, Table S1), despite the simpler architecture of EvoCellNet and significantly fewer parameters compared to ResNet (*∼*6% of the parameters).

**Figure 3:**
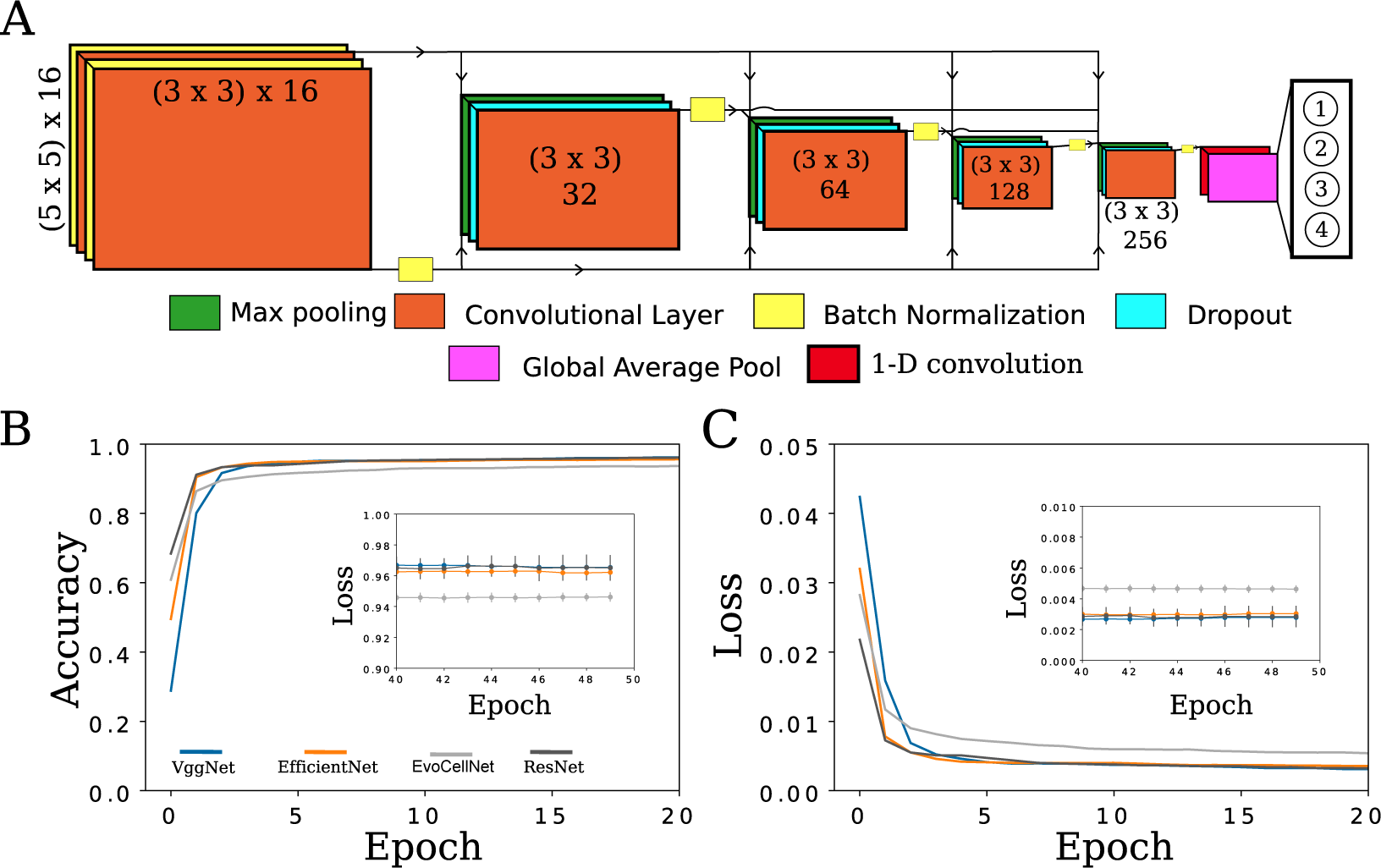
CNN architecture and training metrics. (A) The architecture of our custom convolutional neural network (CNN) is shown. The CNN consists of 6 convolutional blocks and each block consists of multiple color-coded functions: convolution (orange), max pooling (green), batch normalization (yellow), dropout (cyan), global average pool (magenta) and 1D convolution (red). The numeric values indicate filter size (height x width) and the number of filters in the respective layer. The outputs of the classifier (1-4) are fixed. **(B)** The accuracy (left) and loss (right) measure during network training are plotted with epoch number. These are averages of four folds of cross validation. Individual curves are shown in Figure S1. The detailed description of these metrics is described in the *Methods* section. Multiple previously described CNNs- VggNet [30], EfficientNet [31] and ResNet [29] are compared to our network (EvoCellNet).

**Figure 4:**
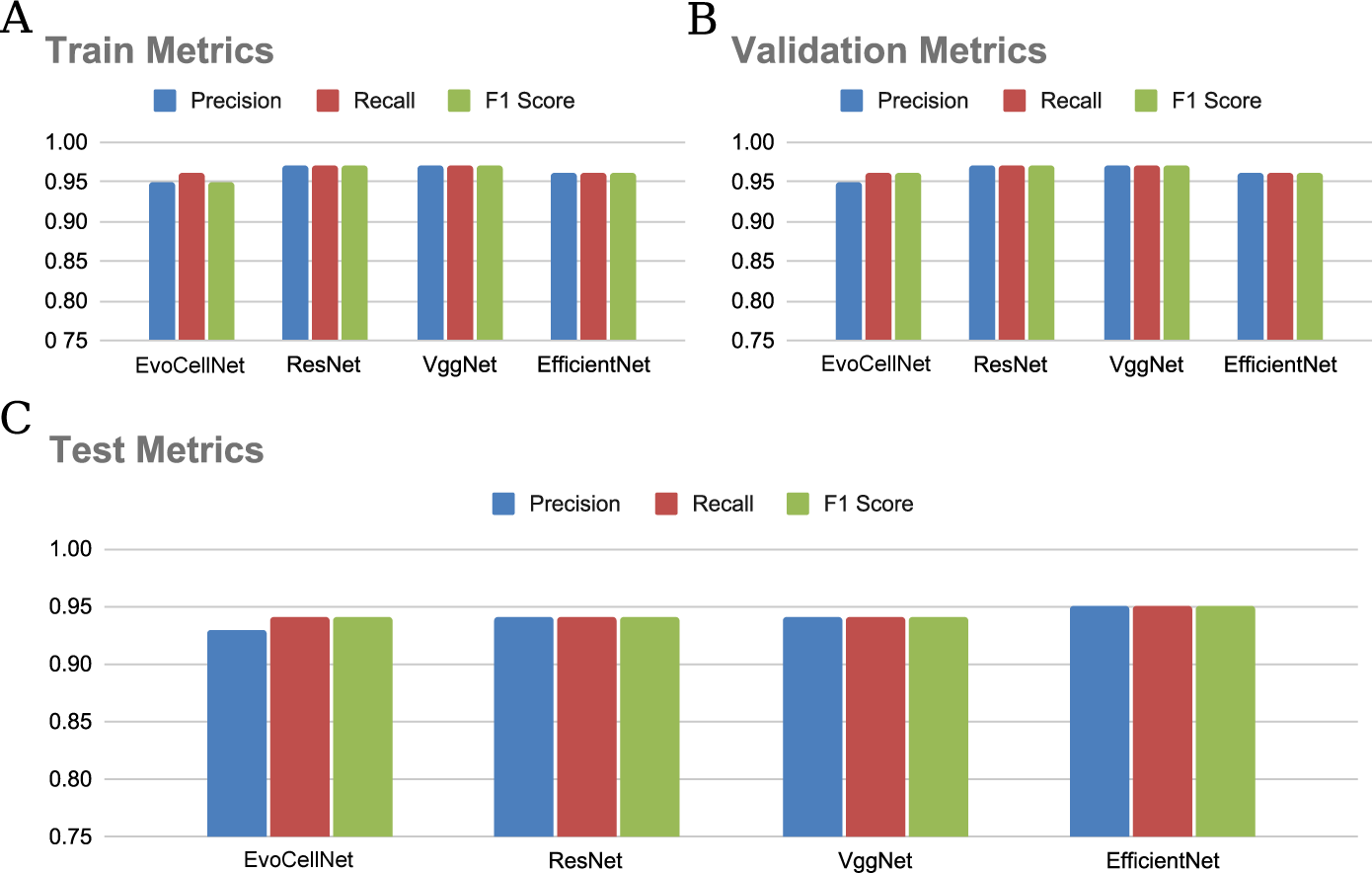
Performance comparison of the trained models in terms of precision, recall and F1 score. Bar plot of Precision, recall and F1 score for the four trained networks: EvoCellNet, ResNet, VggNet and EfficientNet. The bar plots are shown for the (A) training (B) validation and (C) test Dataset.

**Table 2:** Network performace for each stage for validation and test dataset. . The average precision and recall metrics with standard deviation for individual stages for all the four models used in this study. Then mean and standard deviation are calculated across the four folds of the k-fold split for each model.

|  | Pronuclear migration (PM) | Centration rotation (CR) | Spindle displacement (SD) | Cytokinesis (CY) |
| --- | --- | --- | --- | --- |
| **Validation** |  |  |  |  |
| **EvoCellNet** |  |  |  |  |
| Precision | $0.97 \pm 0.010$ | $0.97 \pm 0.004$ | $0.88 \pm 0.012$ | $0.99 \pm 0.003$ |
| Recall | $0.99 \pm 0.004$ | $0.94 \pm 0.013$ | $0.96 \pm 0.007$ | $0.96 \pm 0.009$ |
| **ResNet** |  |  |  |  |
| Precision | $0.99 \pm 0.005$ | $0.97 \pm 0.008$ | $0.94 \pm 0.029$ | $0.98 \pm 0.007$ |
| Recall | $0.98 \pm 0.004$ | $0.97 \pm 0.010$ | $0.93 \pm 0.017$ | $0.98 \pm 0.015$ |
| **VggNet** |  |  |  |  |
| Precision | $0.98 \pm 0.007$ | $0.97 \pm 0.013$ | $0.94 \pm 0.012$ | $0.98 \pm 0.006$ |
| Recall | $0.99 \pm 0.005$ | $0.97 \pm 0.008$ | $0.93 \pm 0.023$ | $0.98 \pm 0.010$ |
| **EfficientNet** |  |  |  |  |
| Precision | $0.98 \pm 0.003$ | $0.97 \pm 0.004$ | $0.93 \pm 0.007$ | $0.98 \pm 0.004$ |
| Recall | $0.98 \pm 0.004$ | $0.96 \pm 0.007$ | $0.93 \pm 0.003$ | $0.98 \pm 0.005$ |
| **Test** |  |  |  |  |
| **EvoCellNet** |  |  |  |  |
| Precision | $0.97 \pm 0.010$ | $0.97 \pm 0.004$ | $0.88 \pm 0.012$ | $0.99 \pm 0.003$ |
| Recall | $0.99 \pm 0.004$ | $0.94 \pm 0.013$ | $0.96 \pm 0.007$ | $0.96 \pm 0.009$ |
| **ResNet** |  |  |  |  |
| Precision | $0.99 \pm 0.005$ | $0.97 \pm 0.008$ | $0.94 \pm 0.029$ | $0.98 \pm 0.007$ |
| Recall | $0.98 \pm 0.004$ | $0.97 \pm 0.010$ | $0.93 \pm 0.017$ | $0.98 \pm 0.015$ |
| **VggNet** |  |  |  |  |
| Precision | $0.98 \pm 0.007$ | $0.97 \pm 0.013$ | $0.94 \pm 0.012$ | $0.98 \pm 0.006$ |
| Recall | $0.99 \pm 0.005$ | $0.97 \pm 0.008$ | $0.93 \pm 0.023$ | $0.98 \pm 0.010$ |
| **EfficientNet** |  |  |  |  |
| Precision | $0.98 \pm 0.003$ | $0.97 \pm 0.004$ | $0.93 \pm 0.007$ | $0.98 \pm 0.004$ |
| Recall | $0.98 \pm 0.004$ | $0.96 \pm 0.007$ | $0.93 \pm 0.003$ | $0.98 \pm 0.005$ |

### The confusion matrix of EvoCellNet is comparable to more complex networks

While overall accuracy and other metrics are comparable we proceeded to examine if the cell cycle stage affected the identification by the networks. The validation dataset of EvoCellNet shows that centration and rotation (CR) is detected correctly with 95.03%, followed by cytokinesis (Cy) with 96.3%, spindle displacement (SD) with 96.68% and pro- nuclear migration (PN) with 98.87% (Figure 5A). For the other networks the score of correct identification of PM is either in the first place for EfficientNet (Figure 5B), or in second place for ResNet (Figure 5C) and VggNet (Figure 5D). At the same time all networks exhibited lower performance in identifying the spindle displacement (SD) stage compared to the other stages in both validation (Figure 5) and test datasets (Figure S2). The misclassification values (off-diagonal) are higher for the CR-SD stage, expected since the transition from the CR stage to the SD stage is marked by the beginning of spindle elongation, making it challenging to precisely identify the exact point of elongation in a frame-wise classification scheme (Figure 5A-D). In contrast, the other two checkpoints, namely the meeting of the two pronuclei and invagination of the cortex, are much easier to identify and can be distinguished at a single time point.

**Figure 5:**
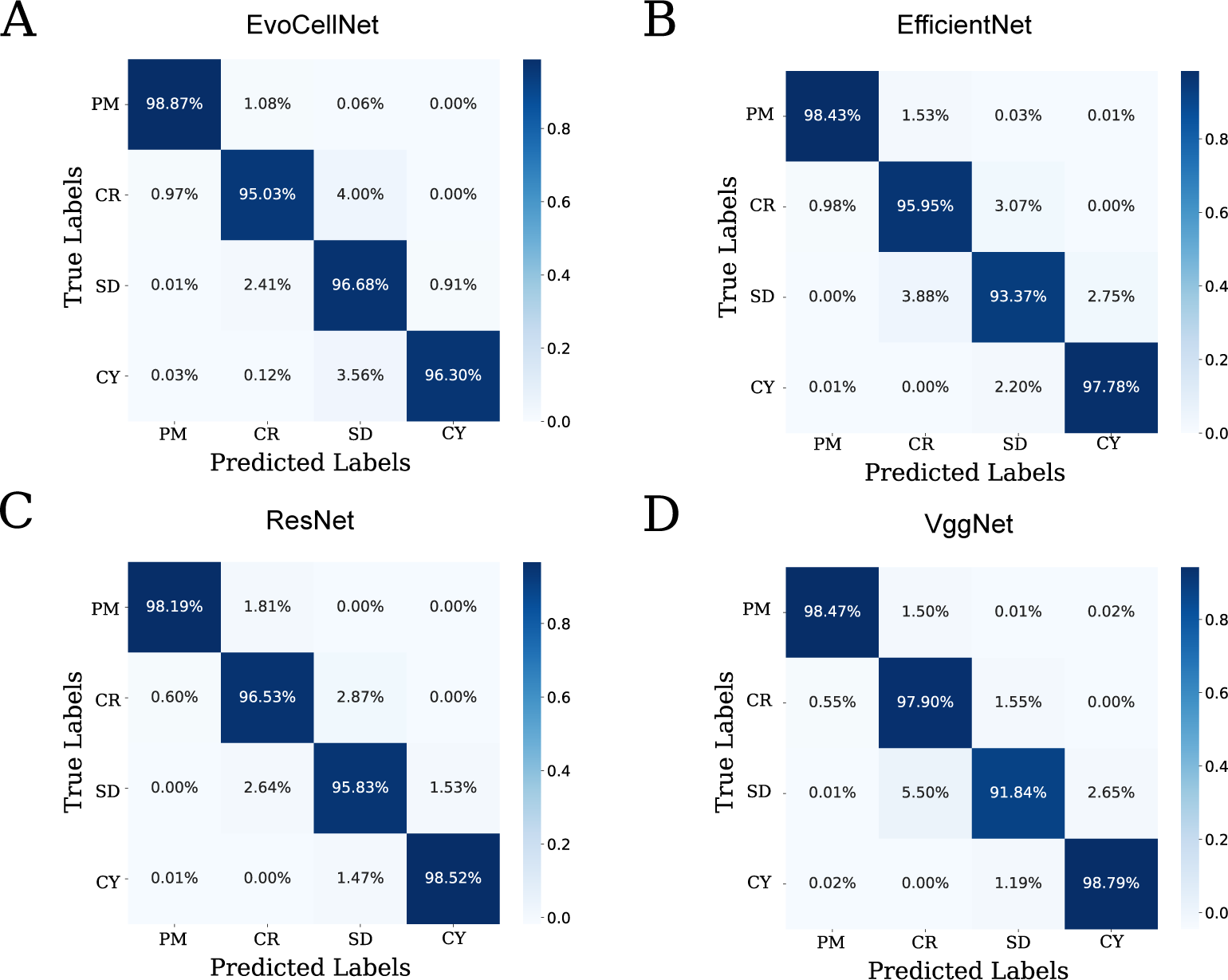
Confusion matrix for the validation set. Confusion matrix with true labels on the y-axis and predicted labels on the x-axis are shown for each model (A) EvoCellNet, (B) EfficientNet, (C) ResNet, (D) VggNet. The matrices are from the first fold of cross validation.

In summary, our results demonstrate the accurate classification performance of the models in identifying the stages of mitotic division. Although the SR stage presented a slightly greater challenge compared to other stages, the overall precision and recall metrics indicate the robustness of the models. To better understand the basis of the classification we attempted to use properties of the network compared to the images.

### EvoCellNet activations identify intracellular structures

The training of CNNs optimizes the weights of convolutional filters so that the classifier results in improved class identification, in this case, based on image features. In order to visualize these features, we use Grad-CAM, developed as a visual explanation of the classification [27], from the last convolutional layer of all models applied to two representative species *C. elegans* and *C. briggsae*. This is one of the approaches used to examine the ‘explainability’ of network outcomes, which could be useful to evaluate biologically relevance and reliability of the method for experts in the field. The rescaled activations represented as a spatial heat-map overlaid on representative DIC images from the time series of each of the stages, and appear to vary between the different networks (Figure 6A-D, Video SV1). In case of EvoCellNet, the activations appear to be co-localized with the visible cytoplasmic structures such as pronuclei, spindle, spindle-poles and parts of the cytokinetic furrow (Figure 6A). In contrast EfficientNet (Figure 6B) and ResNet (Figure 6C) appear to cover larger features of the cells that do not correlate with features used in cell and developmental biology to identify the stages. Finally VggNet appears to result in activations that contain both intracellular features as well as aspects of image-background (Figure 6D). The filters that responded to the transition between uniform and granular intensity profiles seen at the boundaries of the embryo and cytoplasmic structures could explain how EvoCellNet and VggNet result in activations in the pronucleus, spindle region, and cytoplasmic furrow regions. These feature maps demonstrate that some of the trained models use stage-dependent textures to differentiate between classes, similar to the approach taken by an expert analyzing the images.

**Figure 6:**
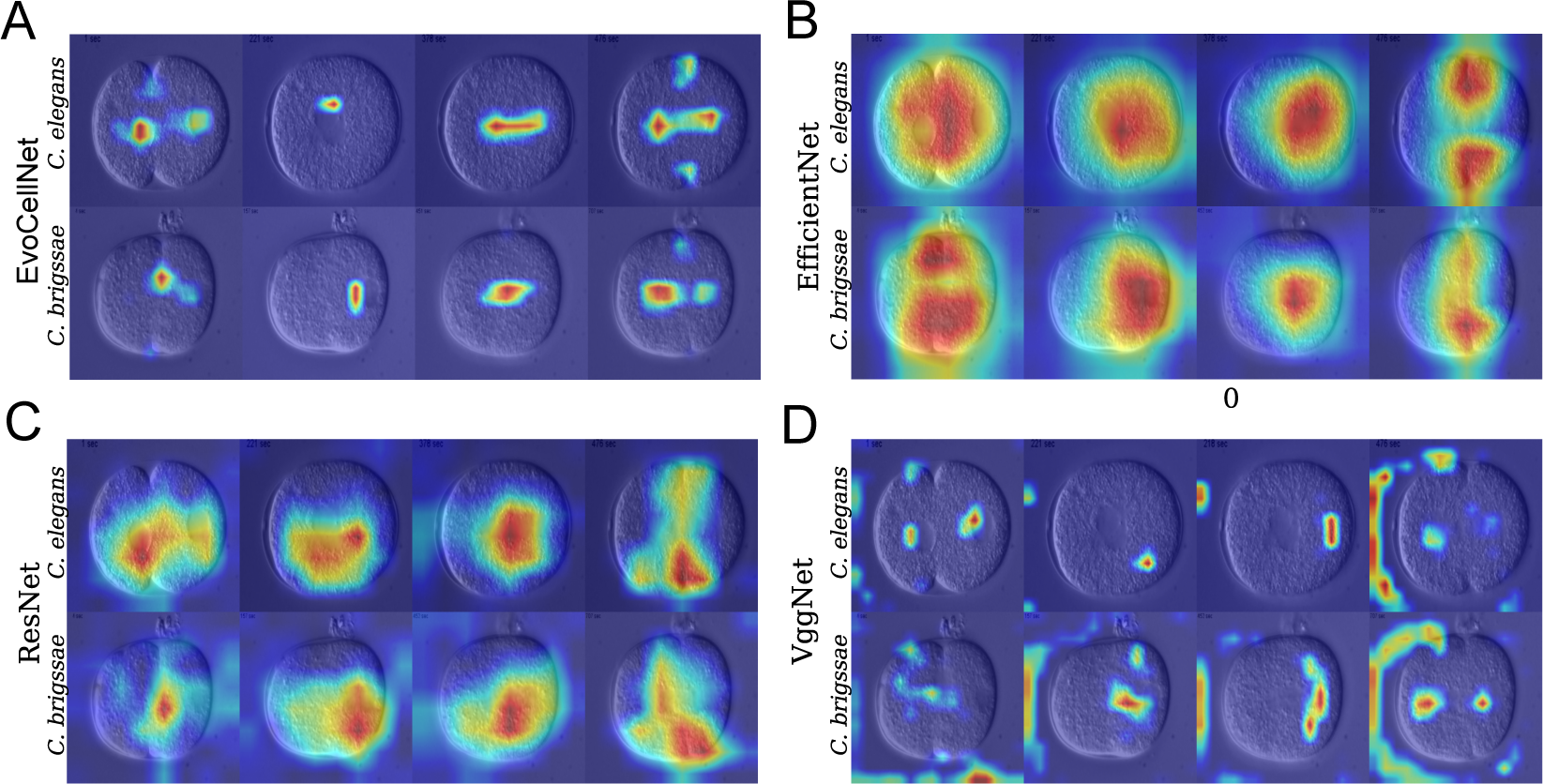
Activations of the last layer or EvoCellNet detect intracellular features. (A-D) A montage of 2 representative species *C. elegans* and *C. briggsae*, with one randomly selected frame from each stage is shown. The images are overlaid with gradient activation values obtained using grad-CAM [27]. Activation values were rescaled to 0 and 255.

With the qualitative assessment of reliability of the networks apparent from Grad-CAM visualization, we proceeded to examine the quantitative error in classification.

### UMAP representation of all networks clusters image time-series cyclically with varied classification errors

In an attempt to obtain quantitative information about the classification and inherent errors, we proceeded to visualize the validation dataset vector in a two-dimensional space with a uniform manifold approximation and projection, UMAP [33]. The UMAP parameters based on Euclidean metrics were optimized to obtain robust results with neighbors = 10, seed = 55, minimal distance = 0.2, number of components = 2. The predefined seed in the UMAP allows us to confirm that our observations are reproducible. The resulting 2D UMAP (in UMAP-1, -2 space) of the validation dataset for *C. elegans* shows each cell division stage (corresponding to color) appears to cluster in a cyclical manner for all networks tested- EvoCellNet, EfficientNet, ResNet and VggNet (Figure 7A). Since the CNNs classify images only in a frame-wise manner, it is interesting that cyclical progression of first embryonic division is captured by the classification without explicit temporal information. These points appear to be concentrated at boundaries between classes in UMAP space, similar to our analysis using the confusion matrix where mis-classifications occur with a higher frequency in adjacent classes (Figure 5). Qualitatively the VggNet appears to have a narrower spread of the misclassified image frames as compared to the other networks, while EvoCellNet exhibits the highest error in classification, as seen in terms of the scatter of misclassified points. The higher error rates between the CR and SD stages for all models are also apparent in the UMAP space. We quantify the errors in four representative species *C. elegans*, *C. briggsae*, *C. remanei* and *C. brenneri* for VggNet (Figure S3A) and EvoCellNet (Figure S4A). The frequency distribution of difference between the individual frames mis-classified either side of a boundary and the manual annotation gives rise to the Δ*T* estimates for the transitions between PM-CR (Δ*T*_1_), CR to SD (Δ*T*_2_) and SD to CY (Δ*T*_3_) that are maximally *±*100 seconds (Figure S3B, S4B). Given that the time series extends over *∼*16 hours, this error is *∼*0.2% of the total time.

**Figure 7:**
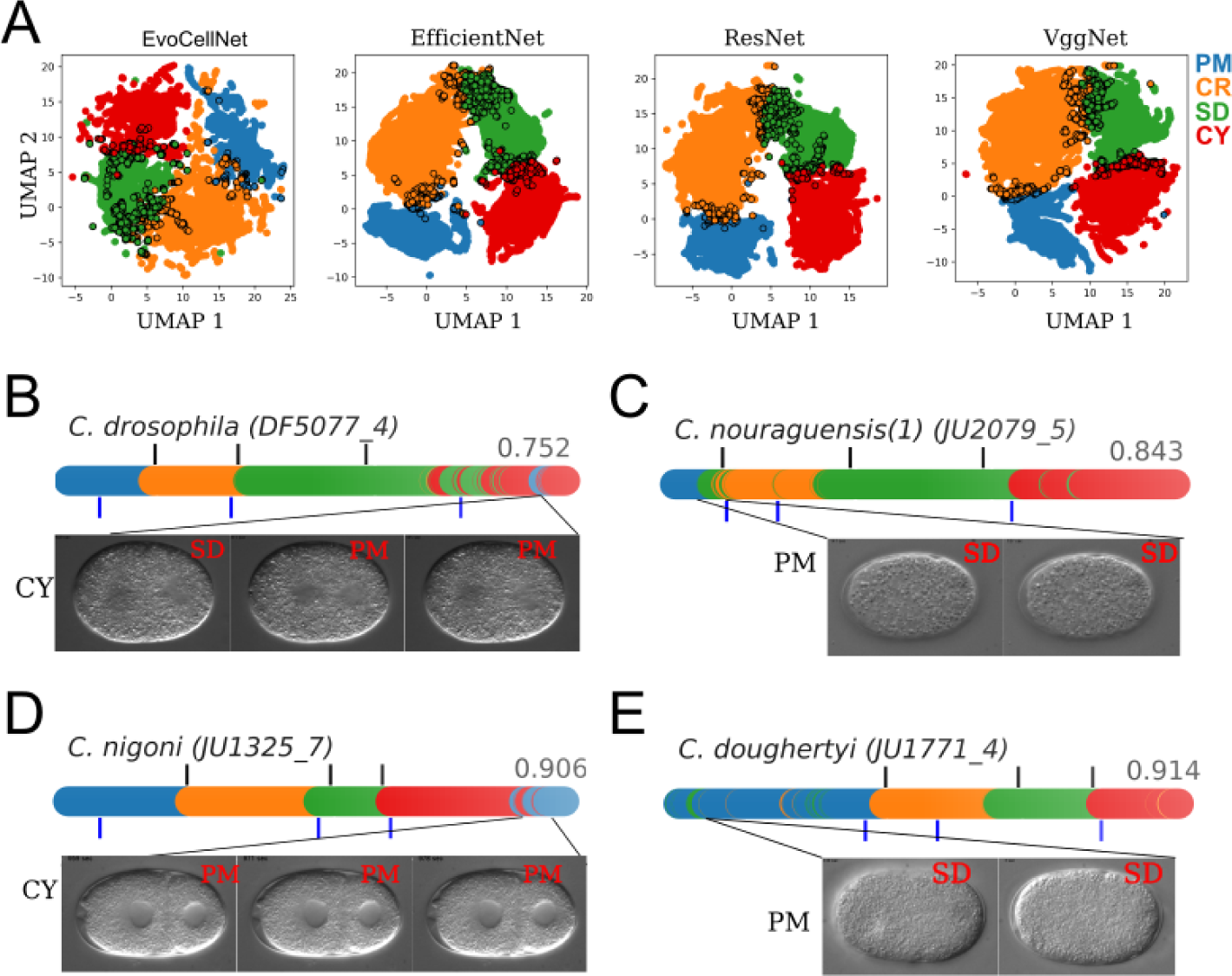
Chronological distribution in UMAP space and the source of misclassifications. (A) All species classifications are represented as a UMAP in 2D based on the feature vector of four classes obtained from the last convolutional layer comparing *(left to right)* EvoCellNet, EfficientNet, ResNet and VggNet. The data points (filled circles) are color coded according to the class, while incorrect classifications are shown as solid circles with black boundaries. The cell division stage to color mapping is as follows PM: blue, CR: orange, SD: green and CY: red. (B-E) Time-series where the CNN classifier performed with less than 92% accuracy are shown in terms of colored plots of frame wise classification output from the ensemble predictor. Black vertical lines mark the manually labelled transition boundaries and blue lines mark the automatically calculated boundaries from SVM (described in detail in the Materials and Methods section). The mis-classified images are shown below each plot with the CNN based classification (red text) and the manual annotation (black text) for (B) *C. drosophila*, (B) *C. nouraguensis*,(D) *C. nigoni* and (E) *C. doughertyi*. The value indicates the classification accuracy between 0 and 1.

The errors in classification misclassified images from EvoCellNet and find that the CNN incorrectly classifies cytoplasmic stage (CY) as pro-nuclear migration stage (PM). This is due to poor visibility of the cytoplasmic furrow. Also, the images contain features that resemble pronulcear migration stage. In the case of (B) *C. nouraguensis* and (D) *C. doughertyi* there are no visible cytoplasmic structures in the plane of acquisition. This results in miss-classification errors (Figure 7B-E)

### Reconstructing the first embryonic cell division stages of ***Caenorhabditis sp.*** from VggNet classifications

With an accurate classification output, we can use the UMAP representation to mark the positions of key cellular events, which were previously manually annotated. By using a linear SVM kernel on the 2D output from UMAP, we can automatically identify the positions of the defined checkpoints between adjacent classes. The absolute difference between the manually annotated time boundary and the automatically found separation boundary (Δ*T_n_*, where n=1-3) shows VggNet has the smallest errors in Δ*T*_1_ and Δ*T*_2_ while Δ*T*_3_ is comparable between VggNet and EfficientNet (Table 3). Thus we proceed to use the combination of VggNet and SVM to reconstruct first embryonic division stages. In order to reconstruct the cyclical nature of cell division, we also plotted a radial centroid with fixed angular period (12*°*) to find the central tendency, initializing the approach to the first time-point in the image time-series. We are now able to represent the UMAP of VggNet classification of the test data as a cycle PM-CR-SD-CY for 4 representative species *C. elegans* (Figure 8A), *C. briggsae* (Figure 8B), *C. remanei* (Figure 8C) and *C. brenneri* (Figure 8D).

**Figure 8:**
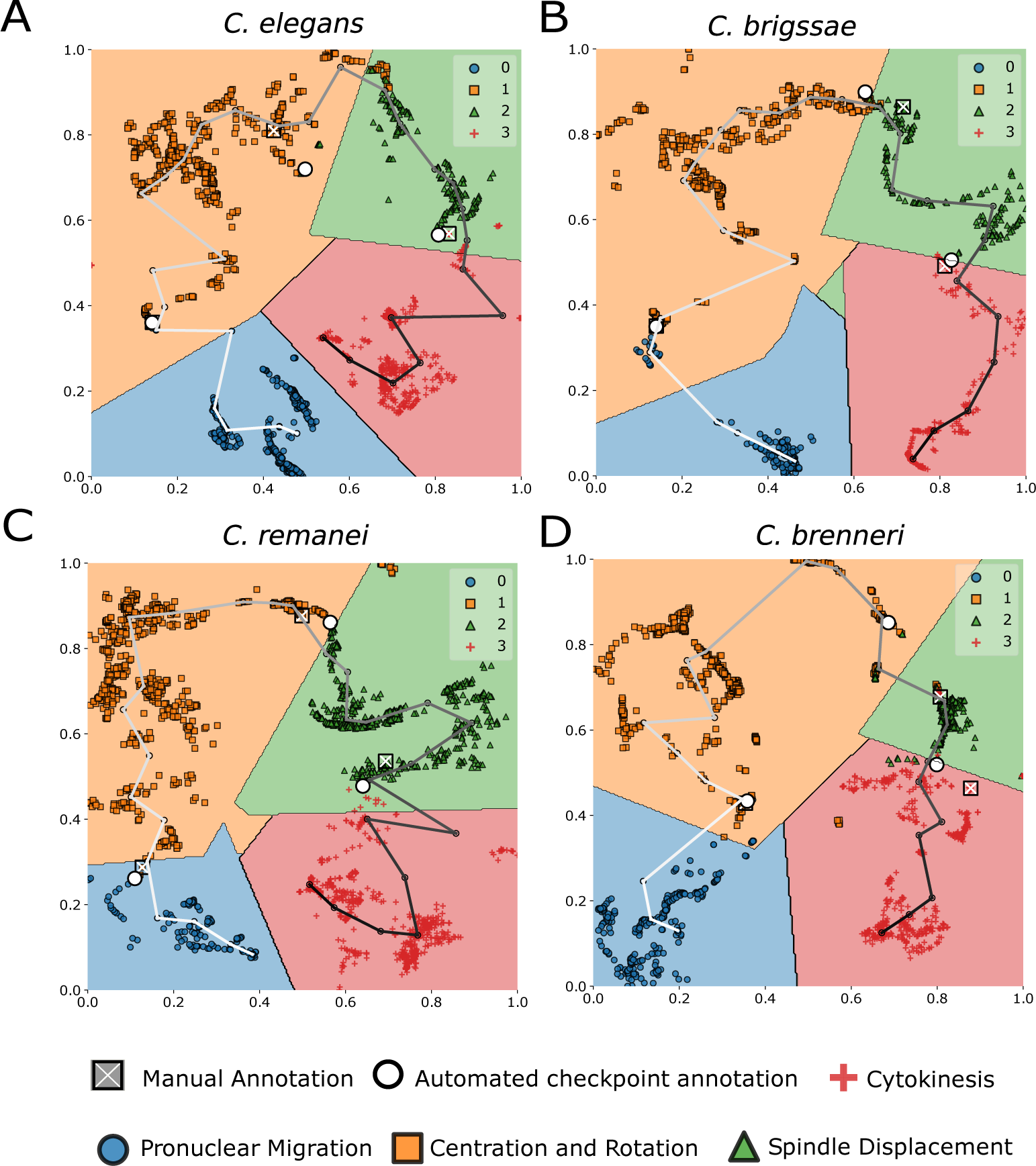
Cell stage boundaries combined with radial density trace in VGGNet classified UMAP space. (A-D) Normalized UMAP representations in (x-axis) UMAP1 and (y-axis) UMAP2 space from VggNet classification of (A) *C. elegans*, (B) *C. briggsae*, (C) *C. remanei*, and (D) *C. brenneri*. The classifier outputs in terms of PM (blue circles), CR (orange squares), SD (green triangle) and CY (+) are used to find boundaries using SVMs as described in the Methods section. The points are used to generate angular density by binning every 12*°* and the radial traces connected geometrically with 0*°* corresponding to the first frame. The line color is a grayscale gradient with the transition of grayscale indicating position relative to start (white) to end (black) of the cycle. The direction set by the transition boundary between PM and CY. x: ground truth of the annotated checkpoint, *◦*: checkpoint from SVM.

**Table 3:**
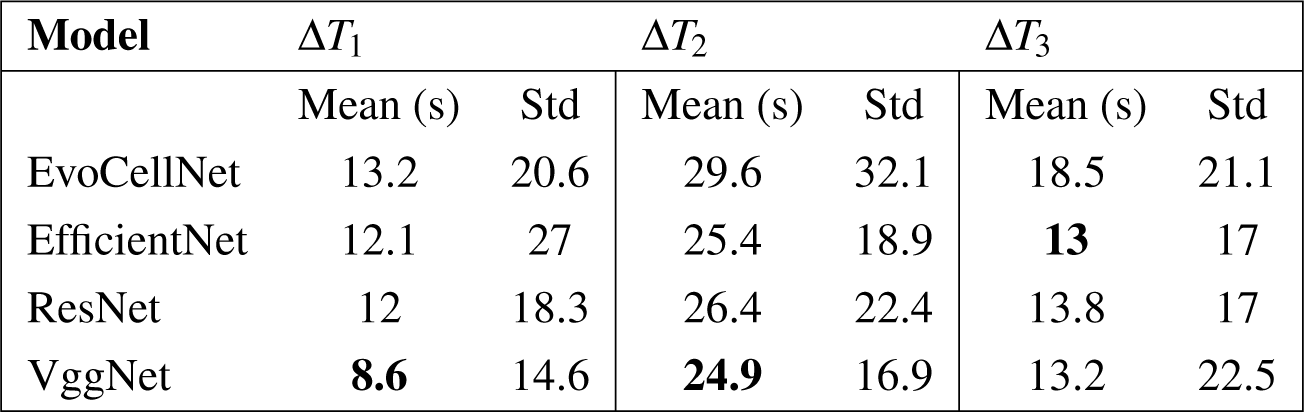
Error in checkpoint prediction by four CNNs. The difference between the manually annotated time of transition and the identification by EvoCellNet, EfficientNet, ResNet and VggNet are expressed for the checkpoints between PM-CR (Δ*T*_1_), CR-SD (Δ*T*_2_) and SD-CY (Δ*T*_3_). The values of the mean error in seconds and standard deviation (Std) are shown for the trained models.

| Model | $\Delta T_1$ | | $\Delta T_2$ | | $\Delta T_3$ | |
| --- | --- | --- | --- | --- | --- | --- |
|  | Mean (s) | Std | Mean (s) | Std | Mean (s) | Std |
| EvoCellNet | 13.2 | 20.6 | 29.6 | 32.1 | 18.5 | 21.1 |
| EfficientNet | 12.1 | 27 | 25.4 | 18.9 | **13** | 17 |
| ResNet | 12 | 18.3 | 26.4 | 22.4 | 13.8 | 17 |
| VggNet | **8.6** | 14.6 | **24.9** | 16.9 | 13.2 | 22.5 |

Thus this approach of CNN based classification allows us to identify cellular transition through male and female pronuclear migration to cytokinesis in multiple *Caenorhabditis sp.* embryos from label-free DIC time series and recovers a cyclical pattern with frame-wise recognition.

## Discussion

Classifying cell stages without the aid of molecular markers is challenging and performed often interactively with qualitative indicators of what might be used as criteria. Here we train 4 different convolutional neural networks and compare their ability to identify the stages of first embryonic division: pronuclear migration (PM), centration and rotation (CR) of the nuclei, spindle displacement (SD) and cytokinesis (CY), from time-series of 21 evolutionarily related species of *Caenorhabditis sp.* previously reported by Valfort et al. [13]. Of these networks, three are state-of-the-art CNNs that have previously been used in image-recognition tasks- VggNet, EfficientNet and ResNet. We also develop a shallow customised network, with comparatively fewer layers, that we refer to as EvoCellNet. Image augmentation and cross-validation schemes are used to balance out the number of images in each category. We find the training of all 4 CNNs to classify the stages of the first embryonic division results in 90% accuracy and comparable training metrics, irrespective of the type of CNN. The confusion matrix of stage-wise comparison of true labels and predictions demonstrates differences between networks with errors at the interface between stages. Using GradCam analysis we find that EvoCellNet appears to surprisingly detect intra-cellular features such as pronuclei, spindle and cytokinetic furrow with no explicit input. The deeper networks appear to detect more global properties of the images with VggNet combining both. We analyze classification errors and them to consist of primarily images at transitions between stages, those with low contrast or morphological similarity with other stages. VggNet scores the highest in terms of test metrics and the outcome on test data is represented in a UMAP. It appears to result in the reconstruction of the cyclical sequence of embryonic division stages, maintaining chronology, despite the lack of any prior information.

In the past classification of microscopy images to monitor sub-cellular changes were attempted using classifiers on cells with molecular markers, but were found to be limited by the image resolution [34]. An approach combining image-segmentation of fluorescent markers with machine learning through multi-class support vector machines (SVMs) could successfully classify cell phenotypes as interphasic and mitotic in RNAi microscopy-screens [35]. The methodological aspects required for a machine learning based proteome screen using microscopy were reviewed, based on feature extraction [36]. Time-series images acquired in high-content screens were classified using a combination of classification hidden Markov models (HMM) [37] for a long time been A segmentation approach of microscopy data to extract feature vectors approach has demonstrated classification of fixed images of cell cycle has typically been identified in every cell type by morphological features and some of it has been captured in work using DNA content, nuclear area and cell crowding to infer cell progression from image based datasets [38]. Indeed Phenotypic classification of cells for drug discovery has demonstrated the use of deep learning by applying them to multi-channel microscopy data of cells treated with compounds [25]. Multi-scale CNNs were used to similar cellular phenotype classification without any low-level image feature extraction with probabilities generated by the network to produce a quantitative and continuous measure of phenotype [39]. More recently label-free data of cell culture using benchtop microscopes was achieved with a ConvNet architecture without supervision with self-label clustering approach to identify cell stages during differentiation [40]. The cell cycle stages G1/S and G2 were also shown to be identified by a deep learning approach based on microscopy data of labelled cell nuclei, Golgi and microtubules, and Grad-CAM analysis was used to confirm the robustness of classification [24]. Our work demonstrates that both the comparative shallow EvoCellNet and the deeper VggNet appear to detect features such as spindle poles and pro-nuclei without the networks having been explicitly provided with any features. This could be used to develop better, interpretable classifiers.

With the Grad-CAM analysis, we demonstrate that the decision-making of our networks is strongly dependent on visible cytoplasmic features. By tuning the network depth, we were able to restrict the gradients only to the most important differences among classes. In EvoCellNet, the final classification is precisely overlaid with cytoplasmic features. This, however, does not mean that other networks’ predictions are not dependent on these features. By reducing the network depth, we were able to increase the focus of convolutional filter parameters only on the cytoplasmic structures of the pronucleus, mitotic spindle, and the cytoplasmic furrow. The deeper networks however appear to learn more global features which lead to their superior performance in segregating the different classes. Interestingly VggNet seems to balance the two tendencies, improved segregation between class clusters in UMAP space and better interpretability in terms of GradCAM activation of intracellular features.

In the future, we could explore the integration of time-series data as input (video recognition) to the deep learning models, allowing for the consideration of temporal dynamics in cell stage recognition. We hope to use more advanced time-series modeling techniques to improve the prediction scheme and the quality of the UMAP trajectory within a class sector. The recognition of cell stages based on cytoplasmic features holds the possibility of automating the comparison of the first mitotic division across diverse nematode embryos. The learned features of the trained networks, both deeper and simpler architectures, can be used to compare spindle mobility differences. The possibility to track or segment cytoplasmic features during the first mitotic division can be extended to include non-*Caenorhabditis* species of the nematode genera for a wider range of comparison of cellular events.

## Materials and Methods

### Image annotation based on cell stage identification

We selected 213 recordings of *Caenorhabditis sp.* embryos from a published dataset [13] which initiate before pronuclear congression. Out of the selected 213 recordings, 44 recordings representing 21 unique *Caenorhabditis* species were kept for testing. The remaining dataset was split into four folds for training and validation (3:1) using stratified *k*-fold cross validation split based on species identity. The distribution of individual frames into four classes is shown in Figure 2(B). Since the database focuses on comparing spindle mobility traits, most recordings start after the spindle has already formed. To avoid over-representation of any one stage during our annotation, we selected only those time series that start before pronuclear migration and fusion. Also, we limited our selection to only *Caenorhabditis* species, which form a major portion (84%) of the database. We then manually annotated each time series based on visible sub-cellular morphology. We marked three sub-cellular events as checkpoints: meeting of the two pronuclei, initiation of spindle elongation, and invagination of the embryo cortex. These checkpoints were used to segregate a given time-series into four stages: pronuclear migration (PM), centration and rotation (CR), spindle elongation and displacement (SD), and cytokinesis (Cy). The nomenclature of these stages is motivated by similar use in previous studies on nematode embryo development [41, 42, 17]. Annotation of these stages was done in ImageJ [43].

### Data augmentation and image pre-processing

Data augmentation is an effective technique to reduce over fitting during network training and improves model generalizability [44]. We applied random image transformations to the input training batch such as vertical and horizontal flips (*probability* = 0.6) and affine and image rotation in range of (0,10) and (0,60) degrees respectively (Figure 2A). All images were resized to (256, 256 px) in height and width and intensity normalized with mean and standard deviation of *µ* = 0.48 and *σ* = 0.15, respectively. The mean and standard deviation were calculated from the entire dataset.

### Training parameters

All networks were optimized using stochastic gradient descent algorithm [45] with a learning rate (*α*) of 1e-2 for 50 epochs and a mini-batch size of 64. The categorical cross entropy loss was minimized during optimization. The network training was done on a V100 NVIDIA (16GB) card. Code implementation and network weights are shared on GitHub (https://github.com/CyCelsLab/nematode-Classification). Users can upload their data and classify it using Google Collab Python Notebook here).

### Evaluation metrics

Descriptive metrics such as accuracy, precision and F1-score were used to evaluate network performance. Accuracy is calculated as:

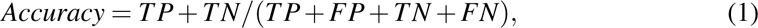

where TP, TN, FP, FN are true positive, true negative, false positive and false negative count respectively. Precision quantifies the proportion of positives that are true from total positives informing us about mis-classification and is described by the expression:

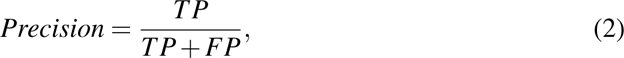

The recall score in turn informs us about the numbers of images that were missed from being classified correctly and is given by:

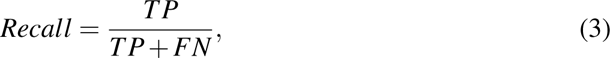

A combined metric that is the harmonic mean of the recall and precision metrics as given by:

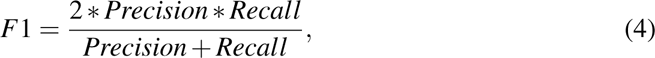

### UMAP representation of network activations

Uniform Manifold Approximation and Projection (UMAP) implementation in python was used [46]. The output feature vector from the trained CNNs was used as the input. Parameters used for UMAP were *n − neighbors*: 10, *seed*: 55, *min − dist*: 0.2, *n − components*: 2, *metric*: “euclidean”. The input embedding were scaled to range [0,1] using *StandardScaler* function provided by *sklearn* library version *0.24* (https://scikit-learn.org/). The *StandardScaler* was fit to the features from the validation dataset for each fold, and is saved for used for future datasets. The UMAP features were rescaled again before plotting. The file used for this is *pseudoTime.py*.

### Transition boundary annotation using SVM

Support vector machine (SVM) algorithm is utilized to determine the transition boundary between adjacent classes. The input to the SVM is a two-dimensional feature vector obtained from UMAP. This feature vector is derived from the prediction outputs of the neural network. The support vectors extracted from the SVM provide potential cellular checkpoints. To filter false positives, we apply a data filtering step where we select time frames that fall within the average support vectors on both sides of the decision boundary. Then, we determine the final transition checkpoint by taking the average of the maximum time frame from the first class and the minimum time frame from the second class. The calculations and analyses were performed using *python ver* 3.8.8, specifically utilizing the SVM algorithm provided by the *sklearn* library version *0.24*. File used for this is *PseudoTime Estimation.ipynb*.

## Acknowledgements

We are grateful to Marie Delattre for valuable discussions about the nematode embryo data. The support and the resources provided by PARAM Brahma Facility under the National Supercomputing Mission, Government of India at the Indian Institute of Science Education and Research, Pune (IISER Pune) are gratefully acknowledged. The project was supported by a grant BT/PR40262/BTIS/137/38/2022 from the Department of Biotechnology, Government of India awarded to CAA.

## Footnotes

## Author contributions

Conceptualization: D.K.; Code and Analysis: D.K.; Writing - original draft: D.K.; Writing - review & editing: D.K., C.A.A.; Funding acquisition: C.A.A.

## Supporting information

### Supporting Tables

**Table S1:** Validation metrics. The average precision, recall, and F1-score metrics with standard deviation for the validation dataset were calculated across the four folds of the k-fold split.

| Network | Precision | Recall | F1 Score |
| --- | --- | --- | --- |
| EvoCellNet | 0.95±0.006 | 0.96±0.004 | 0.96±0.005 |
| ResNet | 0.97±0.006 | 0.97±0.005 | 0.97±0.005 |
| VggNet | 0.97±0.005 | 0.97±0.006 | 0.97±0.005 |
| EfficientNet | 0.96±0.002 | 0.96±0.000 | 0.96±0.001 |

### Supporting Figures

**Figure S1:**
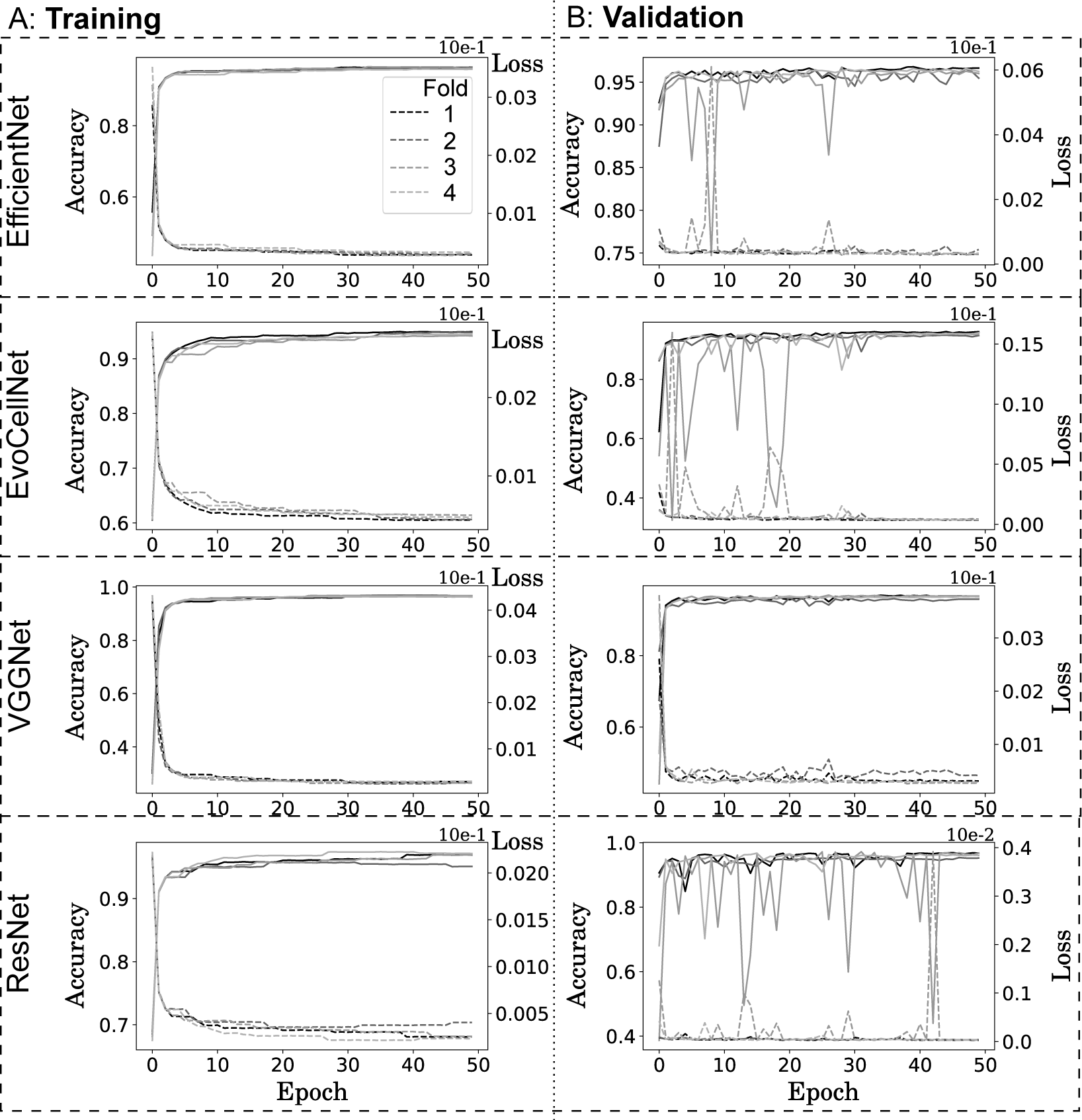
Training and validation metrics for loss and accuracy. The (A) training and (B) validation measured in terms of accuracy (left: left y-axis) and loss (right: right y-axis) for epochs from 0 to 50 for all four folds of training (individual lines) for the EfficientNet, EvoCellNet, VGGNet and ResNet (top to bottom) networks are plotted.

**Figure S2:**
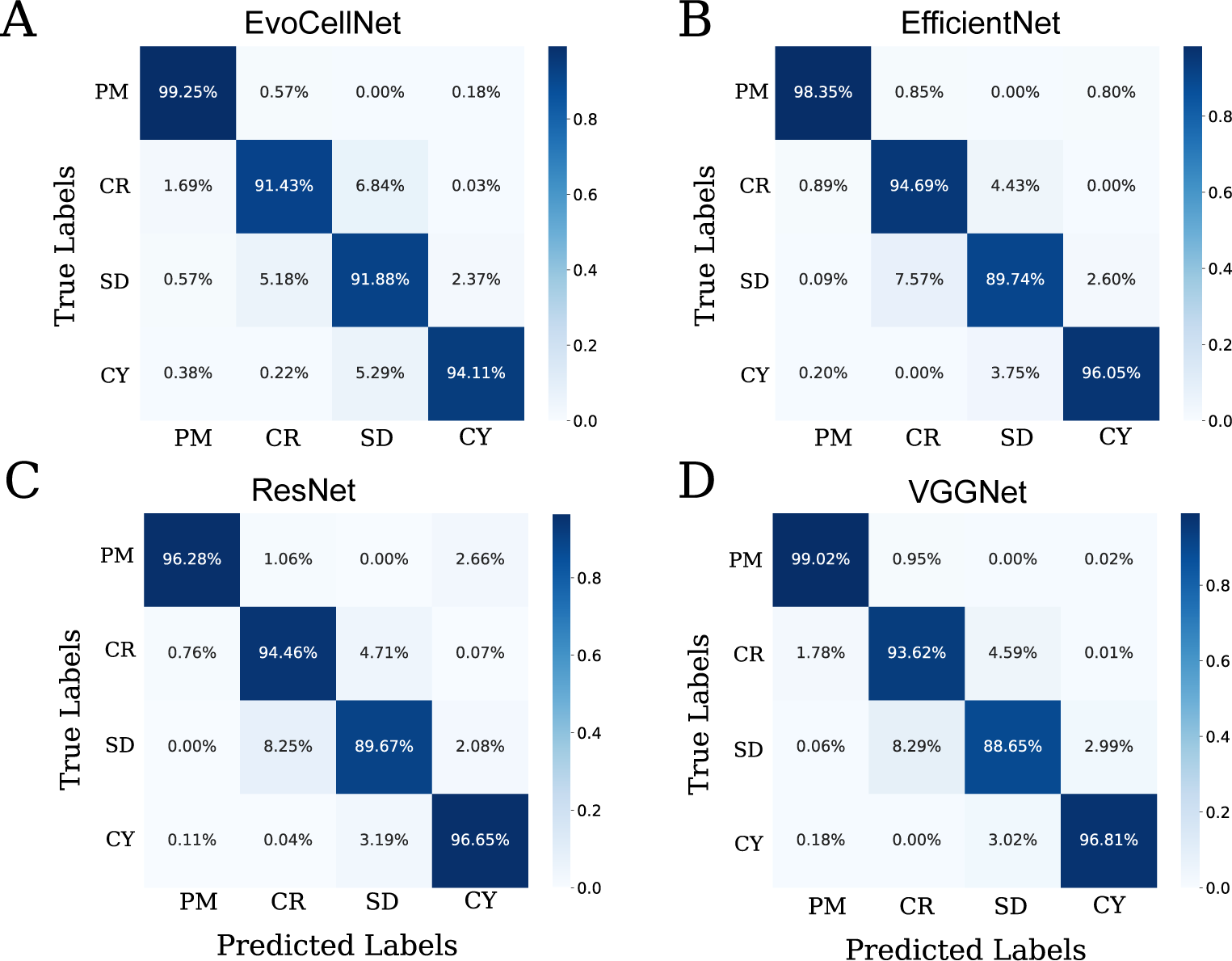
Confusion matrix for the test dataset. Confusion matrix with true labels on the y-axis and predicted labels on the x-axis are shown for each model (A) EvoCellNet, (B) EfficientNet, (C) ResNet, (D) VggNet. The matrices are an calculated from an ensemble of four models trained in each cross validation split.

**Figure S3:**
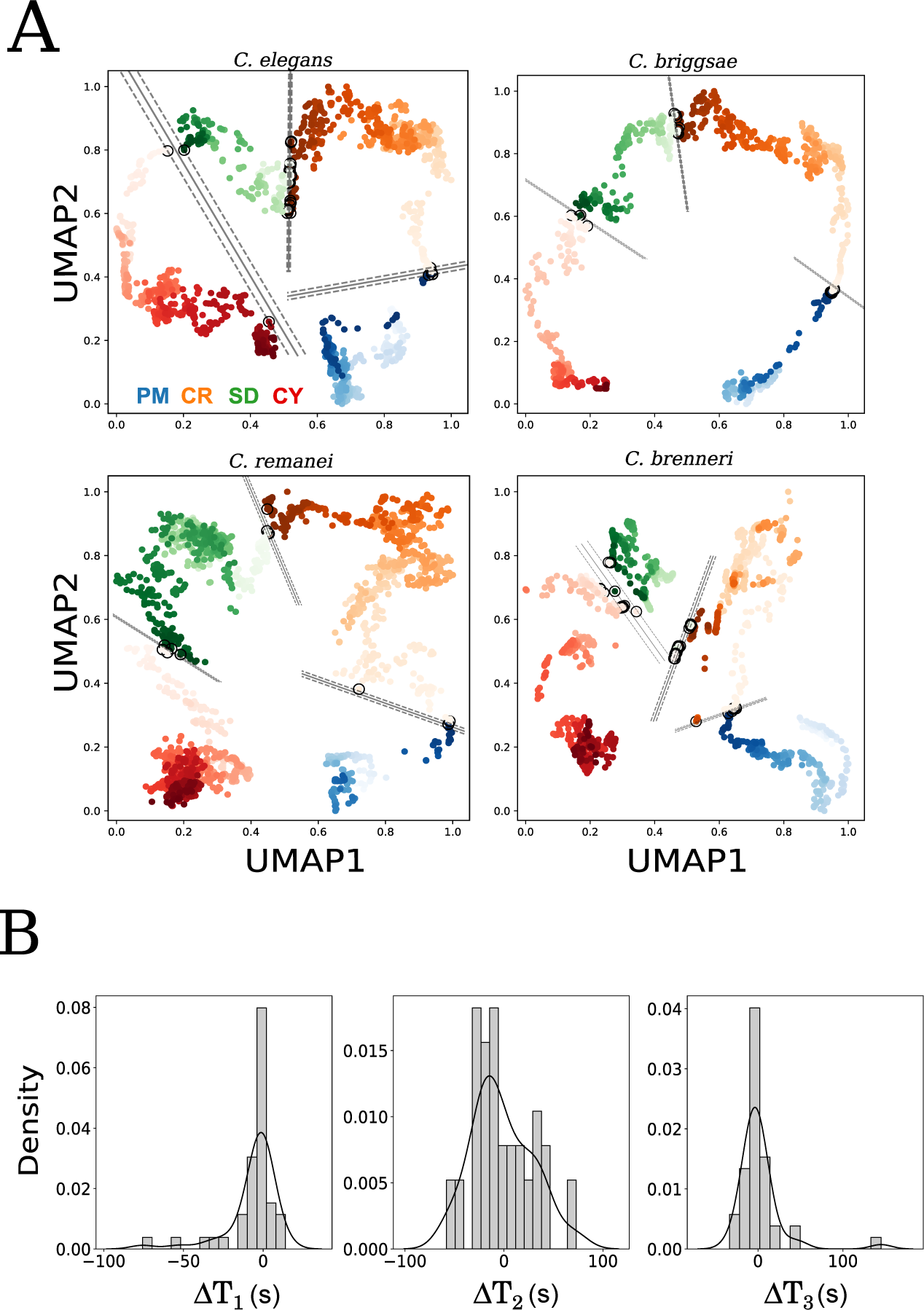
Transition boundary identification by support vector machine (SVM) and error estimation of VGGNet boundaries. (A) The UMAP representation of selected time series from the test dataset for *C. elegans*, *C. briggsae*, *C. remanei*, and *C. brenneri* based on VggNet classifications are plotted. Colors indicate class: PM: blue, CR: orange, SD: green and CY: red. Image-frames (time) are mapped to increasing transparency of the points. Linear SVM kernel is utilized to identify the cell transition boundaries at each adjacent class. The output boundary (solid line) with the 95% confidence interval (dashed line) and support vector (empty circles) are displayed for each UMAP representation demarcating the boundaries. (B) The error in detecting the transition between PM to CR (Δ*T*_1_) CR to SD (Δ*T*_2_) and SD to CY (Δ*T*_3_) in the test dataset is plotted as the histogram of time difference between manually annotated boundary and predicted SVM based on UMAP classifications in seconds. The data is from 21 distinct nematode species related to *Caenorhabditis sp.*.

**Figure S4:**
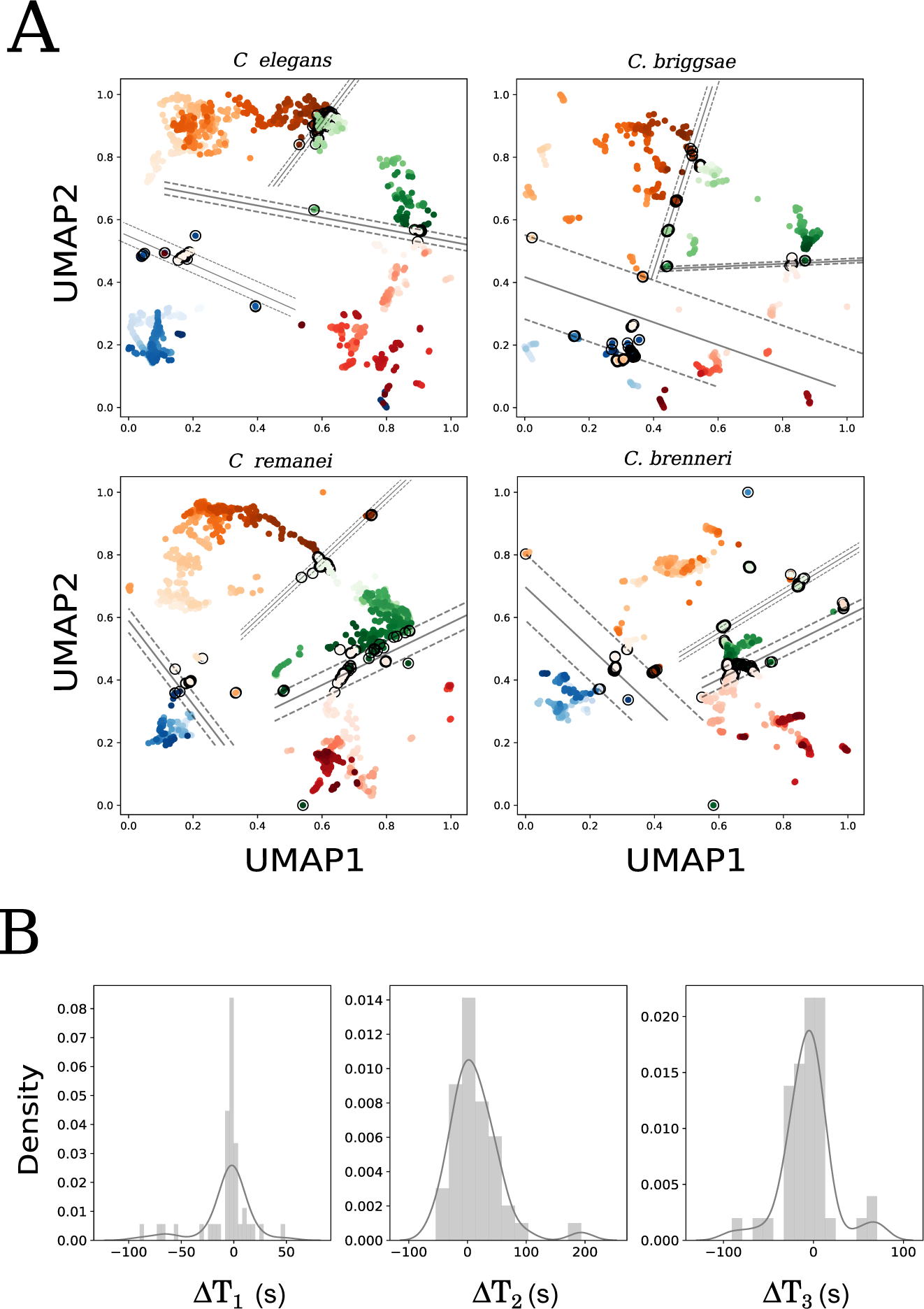
EvoCellNet predictions in UMAP space and the error in estimating transition points between stages. **(A)** The test dataset for *C. elegans*, *C. briggsae*, *C. remanei*, and *C. brenneri* were classified using EvoCellNet and the UMAP representation represents the progression through first embryonic division and a linear SVM kernel was used to mark cell transition boundaries (circles with solid lines). The margins and incorrect classification boundaries are also marked (dashed lines). (B) The difference between manually annotated and SVM predicted boundaries in the test dataset for the transitions between PM to CR (Δ*T*_1_) CR to SD (Δ*T*_2_) and SD to CY (Δ*T*_3_) are plotted as a frequency histogram. The data is from 21 distinct nematode species related to *Caenorhabditis sp.*.

**Video SV1:**
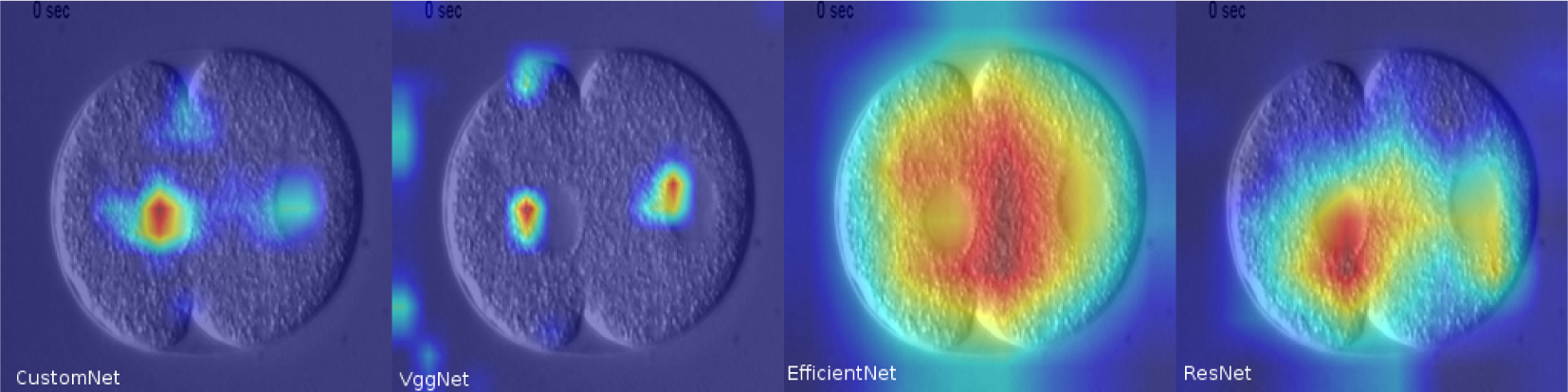
Grad-CAM comparison between four models. Representative filter activations from the 4 models superimposed on the time-series of a *C. elegans* embryo. Activation values are obtained from the last convolutional layer of each model. Colors indicate the activation strength (blue: low, red: high). The models were selected on the basis of lowest validation loss across the four folds of cross-validation.

